# Lysyl oxidase regulates epithelial differentiation and barrier integrity in eosinophilic esophagitis

**DOI:** 10.1101/2023.03.27.534387

**Authors:** Masaru Sasaki, Takeo Hara, Joshua X. Wang, Yusen Zhou, Kanak V. Kennedy, Nicole N. Umeweni, Maiya A. Alston, Zachary C. Spergel, Ritsu Nakagawa, Emily A. Mcmillan, Kelly A. Whelan, Tatiana A. Karakasheva, Kathryn E. Hamilton, Melanie A. Ruffner, Amanda B. Muir

## Abstract

**Background & Aims:** Epithelial disruption in eosinophilic esophagitis (EoE) encompasses both impaired differentiation and diminished barrier integrity. We have shown that lysyl oxidase (LOX), a collagen cross-linking enzyme, is upregulated in the esophageal epithelium in EoE. However, the functional roles of LOX in the esophageal epithelium remains unknown.

**Methods:** We investigated roles for LOX in the human esophageal epithelium using 3-dimensional organoid and air-liquid interface cultures stimulated with interleukin (IL)-13 to recapitulate the EoE inflammatory milieu, followed by single-cell RNA sequencing, quantitative reverse transcription-polymerase chain reaction, western blot, histology, and functional analyses of barrier integrity.

**Results:** Single-cell RNA sequencing analysis on patient-derived organoids revealed that LOX was induced by IL-13 in differentiated cells. LOX-overexpressing organoids demonstrated suppressed basal and upregulated differentiation markers. Additionally, LOX overexpression enhanced junctional protein genes and transepithelial electrical resistance. LOX overexpression restored the impaired differentiation and barrier function, including in the setting of IL-13 stimulation. Transcriptome analyses on LOX-overexpressing organoids identified enriched bone morphogenetic protein (BMP) signaling pathway compared to wild type organoids. Particularly, LOX overexpression increased BMP2 and decreased BMP antagonist follistatin. Finally, we found that BMP2 treatment restored the balance of basal and differentiated cells.

**Conclusions:** Our data support a model whereby LOX exhibits non-canonical roles as a signaling molecule important for epithelial homeostasis in the setting of inflammation via activation of BMP pathway in esophagus. The LOX/BMP axis may be integral in esophageal epithelial differentiation and a promising target for future therapies.

## Introduction

The stratified squamous epithelium of the esophagus is the first line of protection against luminal contents including food, bacteria, and other pathogens. Epithelial barrier disruption as well as impaired epithelial differentiation are universal histologic findings in eosinophilic esophagitis (EoE), a chronic allergic disease that affects children and adults (1,2). Patients with EoE have chronic swallowing issues, vomiting, weight loss and overtime develop esophageal fibrosis and strictures. A better understanding of the perturbations that occur in the epithelial barrier of the esophagus may prevent progression of symptomatology and lifelong esophageal dysfunction.

In our previous publication, we found that lysyl oxidase (LOX), a collagen cross linking enzyme, was increased in the esophageal epithelium of patients with EoE, and to a larger degree in patients with fibrostenosis (3). LOX catalyzes extracellular collagen to form inter- and intramolecular cross-links, thus forming collagen fibers (4). LOX is a requisite for normal tissue structure and integrity with global murine deletion causing perinatal fatality due to aortic aneurysms and pulmonary abnormalities (5,6). In the setting of inflammation, enhanced cross linking within tissue has been shown to promote tissue stiffness in the context of liver fibrosis, cardiovascular disease, and breast cancer (7–9). While its role in perpetuating fibroblast activation and tissue stiffness has been described, little is known about the functional role of LOX outside of extracellular matrix remodeling and tissue stiffness.

LOX has been shown to have non-crosslinking effects in bone, skin, muscle and blood through effects on chemotaxis, gene regulation and differentiation (10–12). It has been shown to be both pro-and anti-tumorigenic, making the organ and the context particularly important in its evaluation (13). In the skin, LOX expression has been demonstrated specifically in differentiated keratinocytes. LOX silencing inhibits keratinocyte differentiation *in vitro* and causes decreased expression of terminal differentiation markers filaggrin (FLG) and keratin 10 (KRT10) (13–15). While it seems to have a role in squamous differentiation, the function of LOX in the esophageal epithelium is unknown. LOX is upregulated in the EoE epithelium, however its role beyond collagen crosslinking in the esophagus has never been explored.

Herein, we sought to determine the role of LOX in the epithelium in the context of EoE inflammation using 3-dimensional (3D) organoid and air-liquid interface cultures. We evaluate the effect of LOX on epithelial differentiation and barrier integrity and describe a novel cytoprotective role for LOX within the inflamed esophageal epithelium.

## Results

### IL-13 induces LOX in the differentiated epithelium in human esophagus

To recapitulate the EoE milieu *in vitro* and to determine the characteristics of LOX in human esophageal epithelium, we performed single-cell RNA sequencing on patient-derived organoids (PDOs) treated with interleukin (IL)-13. Three dimensional esophageal epithelial organoids allow for evaluation of the esophageal epithelial dynamics *in vitro* (16–18). Our prior work demonstrated that stimulation with IL-13, the major effector cytokine in EoE, recapitulates the epithelial reactive changes (such as basal cell hyperplasia) seen in EoE (16–18). PDOs were derived from 3 control subjects (18) and we performed single-cell RNA sequencing on each line in the presence and absence of IL-13. The integrated analysis identified 9 esophageal cell populations in the uniform manifold approximation and projection (UMAP). The 9 clusters were then categorized into 4 groups: quiescent basal, proliferating basal, suprabasal, and superficial, based on the expression of known epithelial makers collagen type VII alpha 1 chain (*COL7A1*), dystonin (*DST*), marker of proliferation Ki-67 (*MKI67*), DNA topoisomerase II alpha (*TOP2A*), tumor protein p63 (*TP63*), involucrin (*IVL*), *FLG*, and desmoglein 1 (*DSG1*) (19,20) (Figure 1A). As expected, high *TP63* expression and low *IVL*, *FLG*, and *DSG1* expression were observed in the basal cluster. We detected *COL7A1* and *DST* transcripts in the quiescent basal, and *MKI67* and *TOP2A* transcripts in the proliferating basal cluster, respectively. In contrast, *IVL*, *FLG*, and *DSG1* were highly expressed in the differentiated cluster group (Figure 1B). We further constructed progression mapping of cell cycle phases. In agreement with expression profiling, the majority of proliferating basal populations and differentiated populations were located in G2/M and G1 cell cycle phase, respectively (Figure 1C).

**Figure 1.**
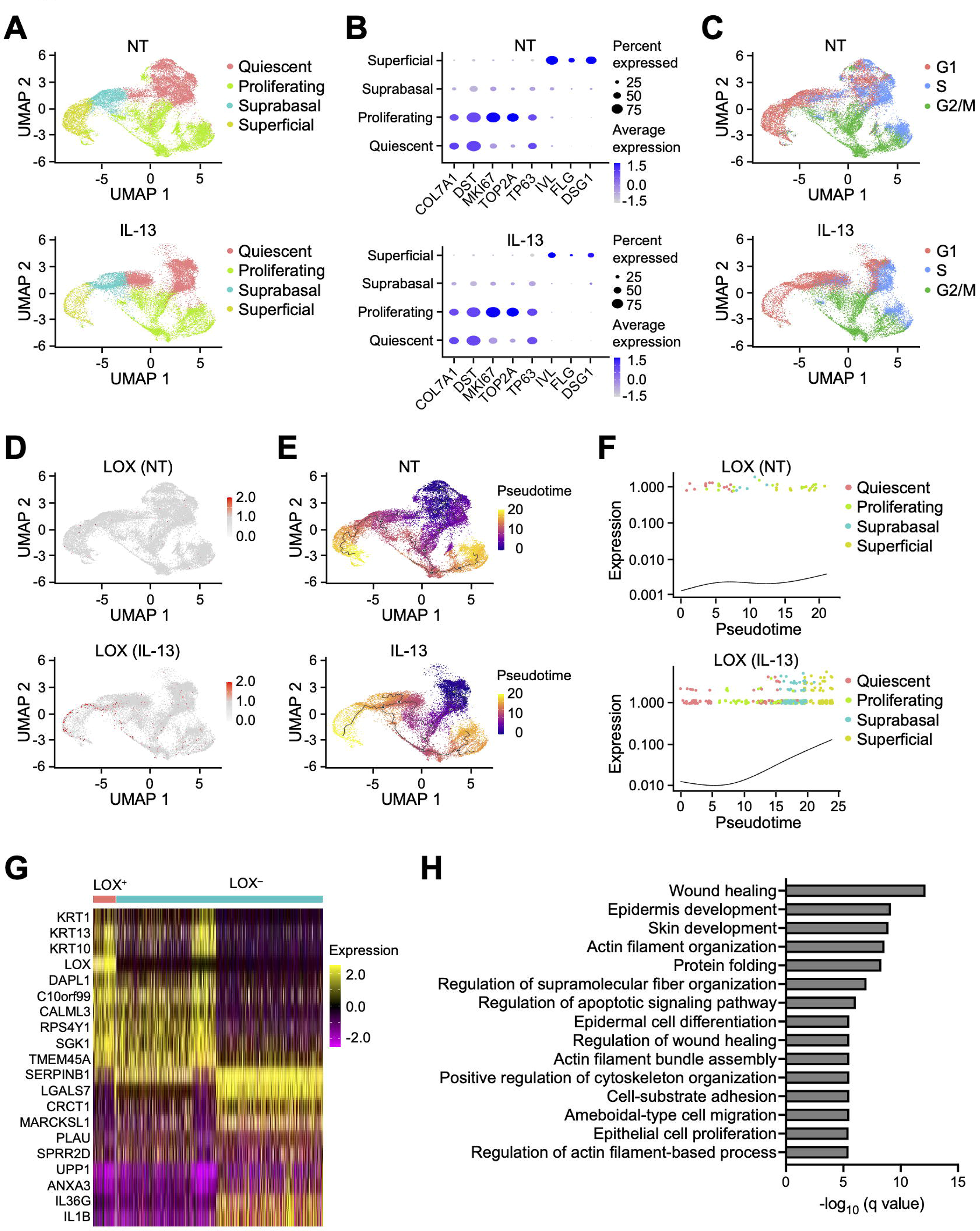

Although *LOX* expression was low in nontreated samples, it was markedly upregulated by IL-13 treatment. Interestingly, cells expressing *LOX* mainly emerged in the differentiated clusters in the UMAP (Figure 1D). Pseudotime analysis was performed to determine developmental relationships between the epithelial populations in human esophagus. Inferred trajectories identified 2 unique cell fates for the quiescent cell cluster in response to IL-13 treatment: (1) toward the proliferating basal cluster and (2) toward the terminally differentiated cluster (Figure 1E). Relative expression values of *LOX* were plotted along pseudotime axis (Figure 1F). The increased expression of *LOX* at the late pseudotime suggests that IL-13 upregulates LOX in differentiated populations within human esophageal epithelium. We further focused on heterogeneity of LOX expressing cells in the differentiated clusters (Figure 1D). Only 10% of the superficial cells were expressing LOX in IL-13-stimulated PDOs. To reveal features of the LOX expressing cells, we performed differentially expression gene (DEG) analysis between LOX expressing (LOX^+^) cells and LOX non-expressing (LOX^-^) cells in the IL-13-treated superficial population. Expression patterns of the top 10 upregulated and downregulated DEGs are shown in Figure 1G. The most highly DEG in the LOX^+^ cells compared to the LOX^-^ cells was keratin 1, followed by keratin 13, and KRT10 which are known as a differentiation marker. Gene Ontology analysis using DEGs demonstrated that heterogeneity of the LOX^+^ and LOX^-^ cells was related to regulation of cell migration, squamous cell differentiation, and cytoskeleton in IL-13-stimulated PDOs (Figure 1H).

### LOX promotes cell differentiation in esophageal epithelium

To investigate the impact of induced LOX in the esophageal epithelium, we overexpressed LOX in the immortalized nontransformed normal human esophageal epithelial cell line (EPC2-hTERT) (3,17,18,21). Green fluorescent protein-transduced (GFP) cells were used as a control. Quantitative reverse transcription-polymerase chain reaction (qRT-PCR) and immunoblotting confirmed increased expression in LOX overexpressing EPC2-hTERT (LOX OE) cells compared to GFP cells in monolayer culture (Figure 2A and 2B).

**Figure 2.**
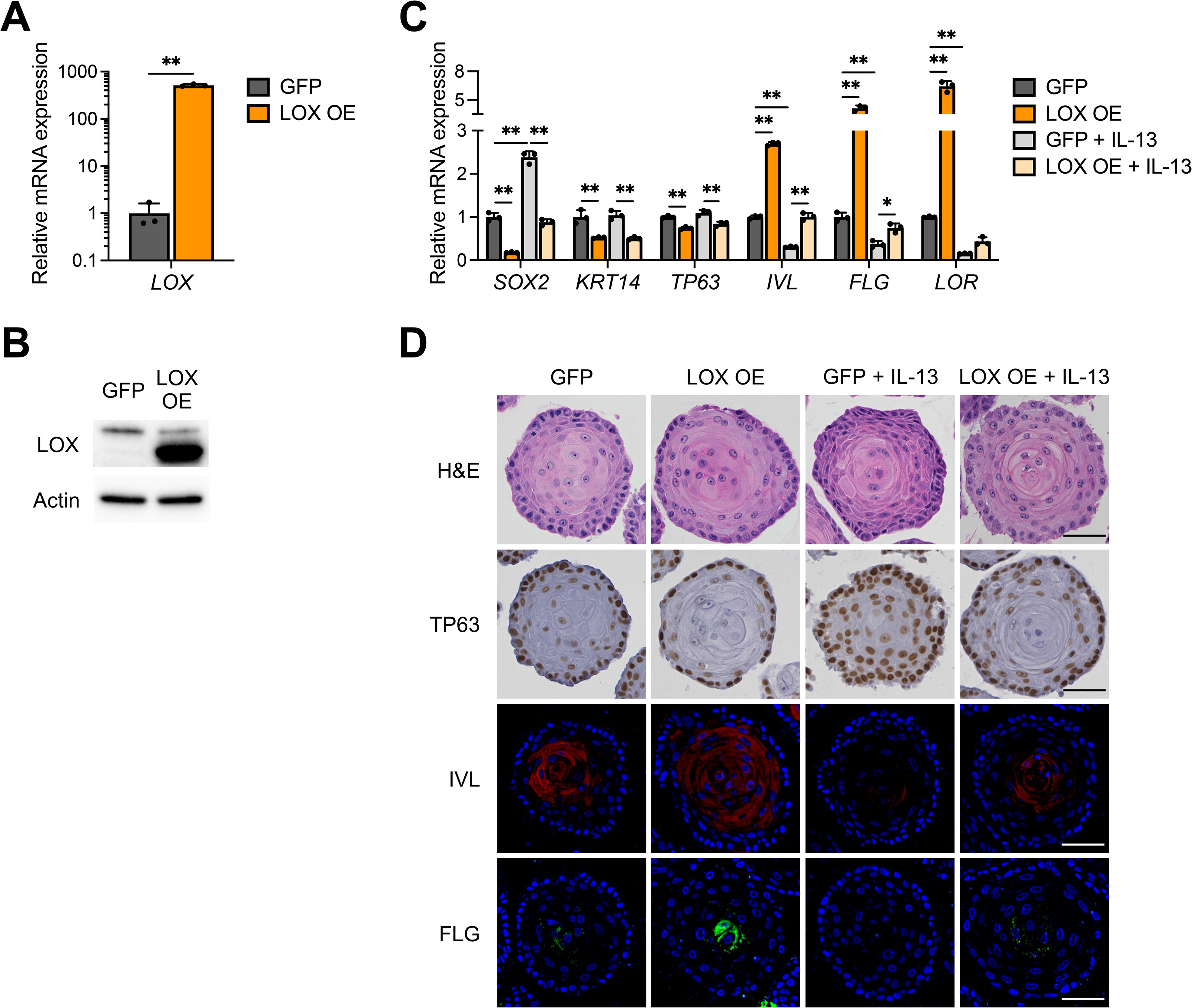
LOX overexpression promotes cell differentiation in esophageal epithelium. (**A**) Quantitative RT-PCR for *LOX* of monolayer-cultured EPC2-hTERT cells overexpressing GFP or LOX (LOX OE) (n = 3). (**B**) Representative images of immunoblot for LOX of the monolayer-cultured GFP and LOX OE cells. (**C** and **D**) Alterations of aberrant LOX expression in EPC2-hTERT organoids. GFP and LOX OE organoids were stimulated with or without IL-13 (10 μg/ml) from day 7 to 11 and then harvested at day 11. (**C**) Quantitative RT-PCR for *SOX2*, *KRT14*, *TP63*, *IVL*, *FLG*, and *LOR* of the GFP and LOX OE organoids (n = 3). (**D**) Representative images of hematoxylin and eosin (H&E) staining, immunohistochemistry for TP63, and immunofluorescence staining for IVL (red) and FLG (green) of the GFP and LOX organoids. DAPI (blue). Scale bar, 50 μm. Data are representative of three independent experiments and expressed as means ± SDs. Two-tailed Student’s t-test (**A**) and one-way analysis of variance (**C**) were performed for statistical analyses. \**P* < 0.05, \*\**P* < 0.01.

We next evaluated 3D organoid cultures (16,17) stimulated with IL-13 (22,23). Ectopic LOX expression resulted in decreased expression of basal cell marker genes SRY-box transcription factor 2 (*SOX2*), keratin 14 (*KRT14*), and *TP63*, and increased expression of differentiation marker genes *IVL*, *FLG*, and loricrin (*LOR*), in both nontreated and IL-13-treated organoids (Figure 2C). We also assessed organoid morphology. Hematoxylin and eosin stain (H&E) staining revealed that LOX OE organoids had advanced inner core hyperkeratosis compared to GFP organoids. IL-13-treated GFP organoids had expansion of the basal cell population, as seen in EoE (1), with thickening of the outer basaloid layer. However, this effect was attenuated in IL-13-treated LOX OE organoids (Figure 2D). Immunohistochemistry and immunofluorescence staining demonstrated elevation of TP63 and depression of IVL and FLG levels by IL-13 in GFP organoids. On the other hand, LOX OE organoids demonstrated reduced expression of TP63 and enhanced expression of IVL and FLG in both untreated and IL-13 conditions compared to GFP organoids (Figure 2D). These findings support the conclusion that LOX partially mitigates the disrupted cellular gradient caused by IL-13 stimulus.

Since only basaloid cells (and not terminally differentiated cells) are capable of forming organoids, we assessed organoid formation rate (OFR) to confirm whether LOX OE cell has reduced OFR due to enhanced differentiation. Although OFR did not significantly change after seeding from 2D basaloid-monolayers into organoids (P0), it was significantly decreased in passaged LOX OE organoids (P1), suggesting that LOX OE organoids comprise more differentiated cells than GFP organoids (Figure 3A and 3B). Taken together, these results confirm that overexpressing LOX promotes epithelial differentiation.

**Figure 3.**
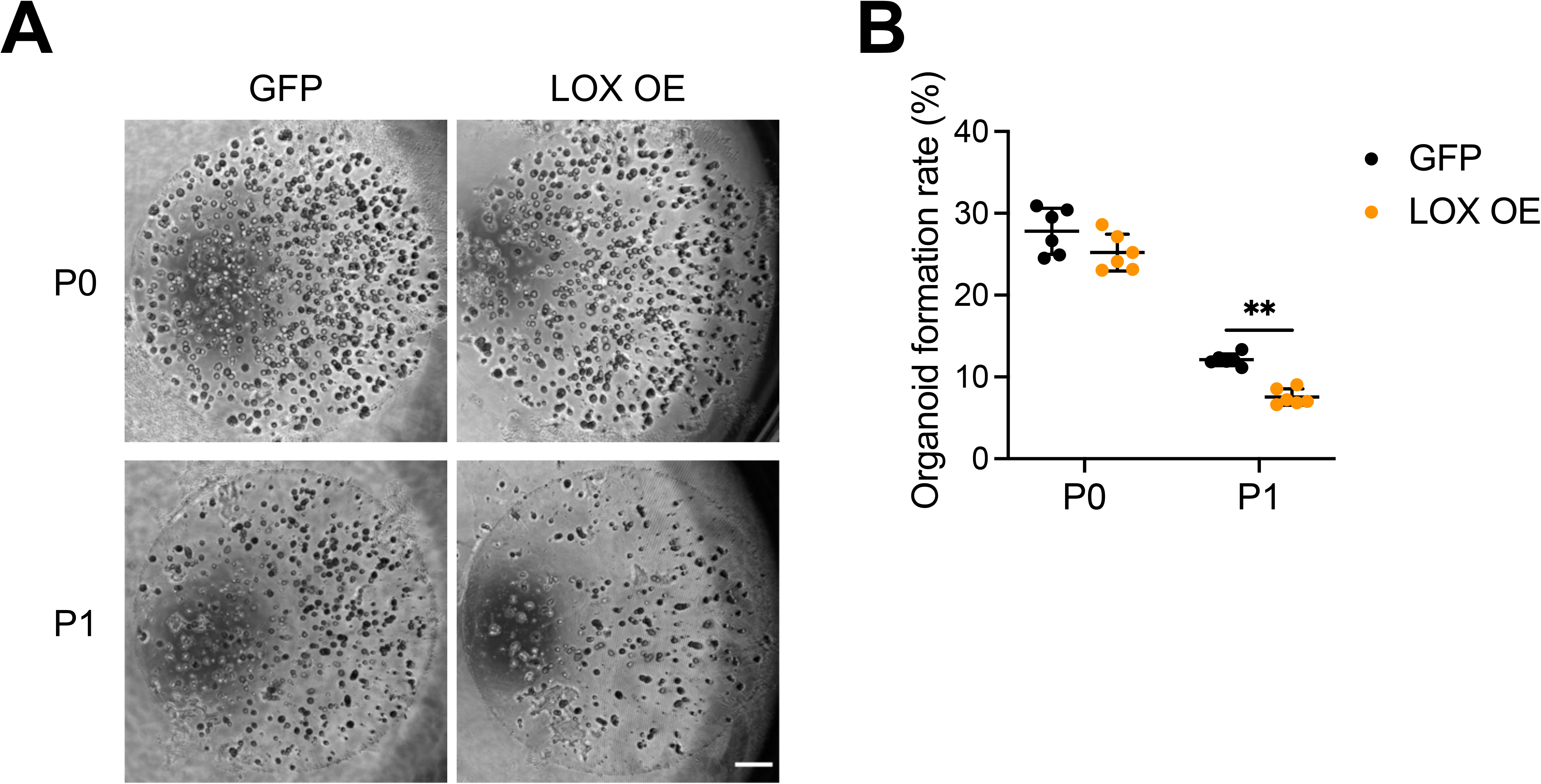
LOX overexpression attenuates organoid formation capacity. (**A**) Representative phase contrast images of EPC2-hTERT organoids overexpressing GFP or LOX (LOX OE) at day 11. Organoid formation rate (OFR) was assessed at day 11 (P0) and they were then passaged. OFR was assessed again at day 11 (P1). Scale bar, 50 μm. (**B**) OFR was defined as the number of organoids (≥ 50 μm) divided by the total seeded cells. Data are representative of three independent experiments and expressed as means ± SDs (n = 6). Two-tailed Student’s t-test was performed for statistical analyses. \*\**P* < 0.01.

### LOX improves barrier integrity in esophageal epithelium

Disturbed squamous cell differentiation alters epithelial barrier integrity (24,25). This disruption is crucial in the pathogenesis of EoE (26–28). Thus, we sought to elucidate if LOX is implicated in barrier regulation. DSG1 and desmocollin-1 (DSC1), members of the desmosomal cadherin family, are required for adhesive intercellular junctions and maintenance of epithelial homeostasis (29,30). Interestingly, while IL-13 treatment decreased transcript levels of *DSG1* and *DSC1* in organoids, LOX overexpression increased their expression at baseline and partially rescued the effect of IL-13 (Figure 4A).

**Figure 4.**
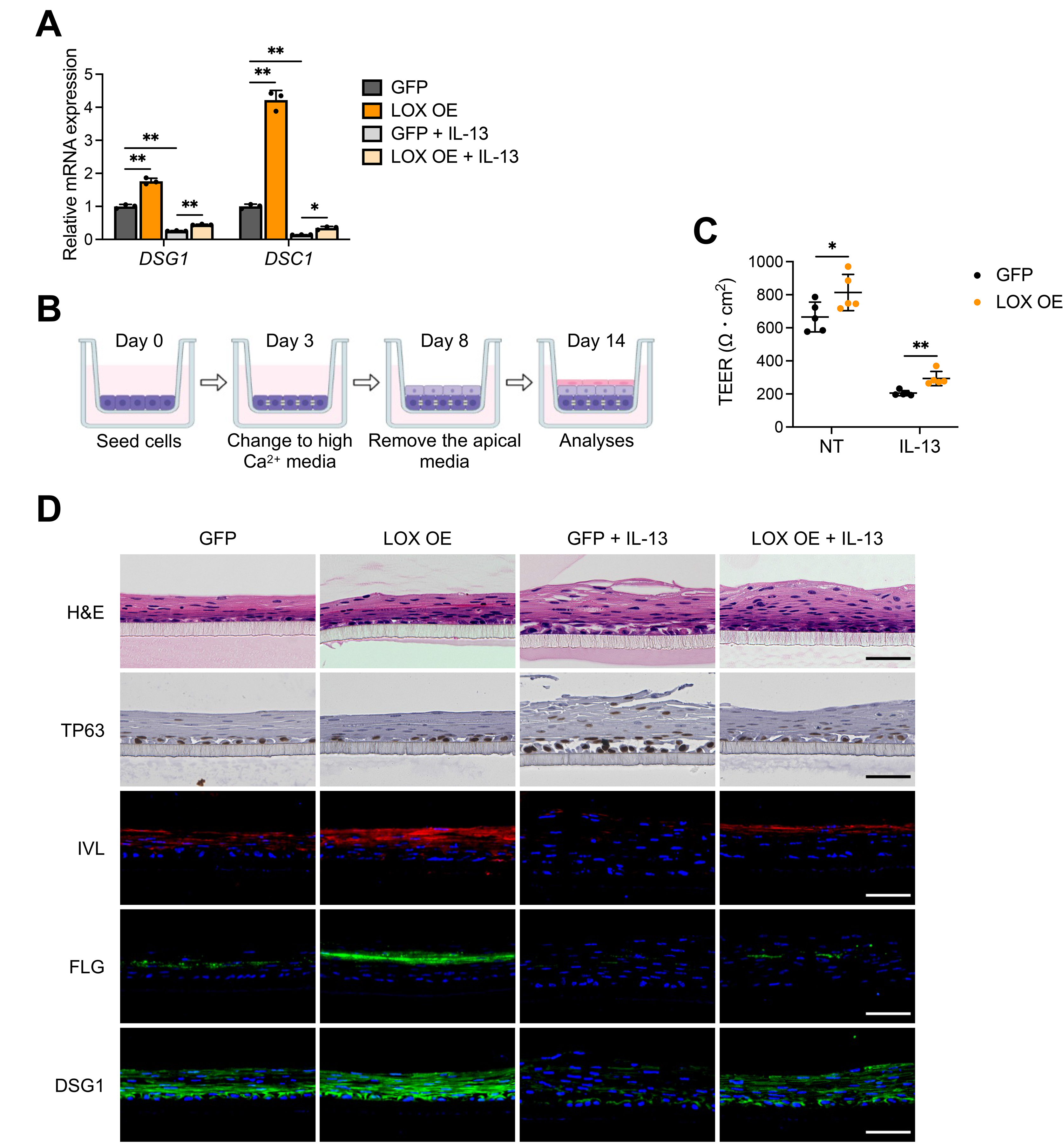
LOX overexpression improves epithelial barrier integrity. (**A**) Quantitative RT-PCR for *DSG1* and *DSC1* of EPC2-hTERT organoids overexpressing GFP or LOX (LOX OE). GFP and LOX OE organoids were stimulated with or without IL-13 (10 μg/ml) from day 7 to 11 and then harvested at day 11 (n = 3). Data are representative of three independent experiments. (**B**) Schematic of air-liquid interface (ALI) model. GFP and LOX OE EPC2-hTERT cells were cultured in low-calcium (0.09 mM Ca2+) media for 3 days, followed by high-calcium media (1.8 mM Ca2+) for 5 days, and then brought to ALI at day 8. ALI-cultured cells were stimulated with or without IL-13 (10 μg/ml) from day 9 to 14. (**C**) Transepithelial electrical resistance (TEER, Ω * cm2) in the GFP and LOX OE EPC2-hTERT ALI-cultures (n = 5). (**D**) Representative images of hematoxylin and eosin (H&E), immunohistochemistry for TP63, and immunofluorescence staining for IVL (red), FLG (green), and DSG1 (green) of the GFP and LOX OE EPC2-hTERT ALI-cultures. DAPI (blue). Scale bar, 50 μm. Data are representative of two independent experiments and expressed as means ± SDs. One-way analysis of variance (**A**) and two-tailed Student’s t-test (**C**) were performed for statistical analyses. \**P* < 0.05, \*\**P* < 0.01. NT, nontreated.

To directly assess the effect of LOX on barrier function, we used the ALI system which mimics an *in vivo* epithelial environment and measured transepithelial electrical resistance (TEER) (31) (Figure 4B). IL-13 treatment increased barrier permeability in the epithelium as measured by TEER. Conversely, LOX overexpression improved the IL-13-induced barrier deficiency by 1.4-fold compared to GFP cells (Figure 4C). H&E staining of GFP cultures also showed impaired squamous stratification in response to IL-13. However, LOX overexpression counteracted the disruption caused by IL-13. We performed staining for TP63, IVL, FLG, and DSG1 in ALI cultures. As seen in organoids, although IL-13 stimulation prevented differentiation and barrier-related protein expression in the epithelium, LOX overexpression enhanced IVL, FLG and DSG1 in both untreated and IL-13 stimulated cultures. Collectively, these data show that LOX promotes normal cell differentiation and supports epithelial integrity including in the setting of IL-13 stimulation.

### Transcriptome profiling reveals pathways associated with LOX expression in esophageal epithelium

To understand the mechanism by which LOX regulates epithelial integrity, we performed RNA sequencing on LOX OE organoids. DEG analysis using DESeq2 (32) identified 2446 upregulated (*P* < 0.05 and log_2_(fold change) ≥ 0.585) and 2923 downregulated (*P* < 0.05 and log_2_(fold change) ≤ -0.585) genes in LOX OE organoids compared to GFP organoids (Figure 5A and 5B). Gene Ontology analysis of DEGs revealed that cell differentiation and keratinization-related terms were enriched in LOX OE organoids (Figure 5C). By contrast, cell proliferation-related terms were decreased (Figure 5D). Furthermore, Gene Set Enrichment Analysis (GSEA) on the Pathway Interaction Database (PID) (33) revealed that gene signatures associated with bone morphogenetic protein (BMP), transforming growth factor-β (TGFβ), and WNT pathways, were significantly enriched in LOX OE organoids (false discovery rate; FDR < 0.25; Figure 5E). These results are consistent with our previous results showing a role for LOX in modulating esophageal epithelial differentiation.

**Figure 5.**
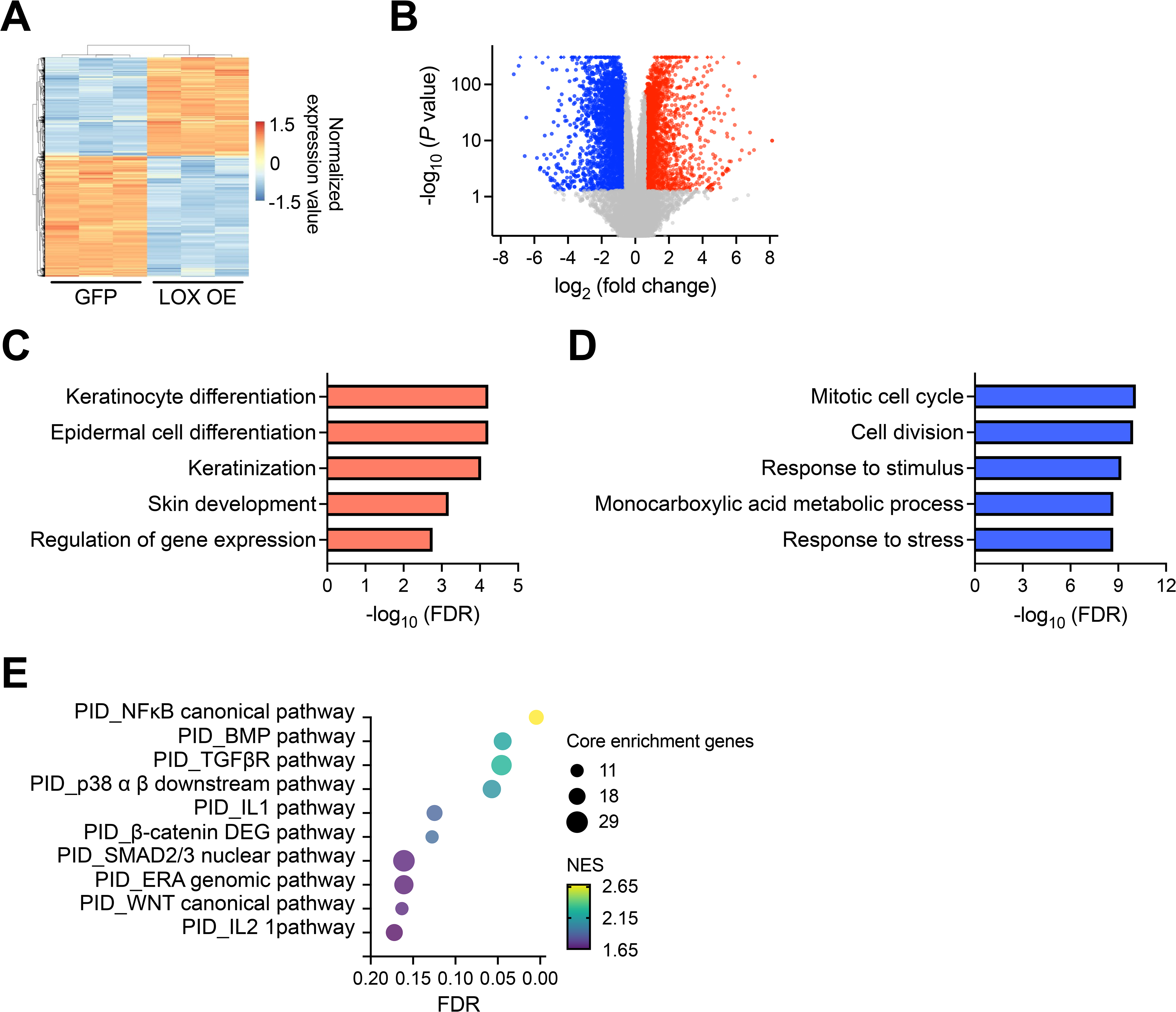
Transcriptome analysis identifies the functional roles of LOX in esophageal epithelium. (**A**) Heatmap and (**B**) volcano plot of differentially expressed genes (DEGs) based on RNA sequencing data from EPC2-hTERT organoids overexpressing GFP or LOX (LOX OE). GFP and LOX OE organoids were cultured for 11 days and subjected to RNA sequencing analysis. Upregulated and downregulated DEGs in LOX OE organoids are shown with red and blue, respectively. (**C** and **D**) Top 5 enriched (**C**) and depleted (**D**) terms in LOX OE organoids based on Gene Ontology analysis. (**E**) Gene Set Enrichment Analysis based on the Pathway Interaction Database (PID). Top 10 enriched pathways in LOX OE organoids are shown. Dot size and color represent the number of core enrichment genes and normalized enrichment score (NES) in the pathway, respectively. FDR, false discovery rate.

### BMP signaling pathway is activated in LOX overexpressing organoids

Recent studies demonstrated that BMP pathway is essential for esophageal progenitor cell differentiation and disrupted BMP signaling results in basal cell hyperplasia in EoE (34,35). Therefore, we postulated that BMP signaling pathway could be integral in LOX-supported differentiation (Figure 6A). We investigated BMP ligands and BMP receptors (BMPRs) from the PID gene sets presented in Figure 5E and 6A. *BMP2*, *BMP6*, *BMPR1B*, and *BMPR2* were significantly elevated in LOX OE organoids compared to GFP organoids (by 2.5-fold, 18.3-fold, 4.8-fold, and 1.2-fold, respectively). Intriguingly, we found that the BMP antagonist follistatin (*FST*) was significantly decreased by 0.36-fold in LOX OE organoids (Figure 6B).

**Figure 6.**
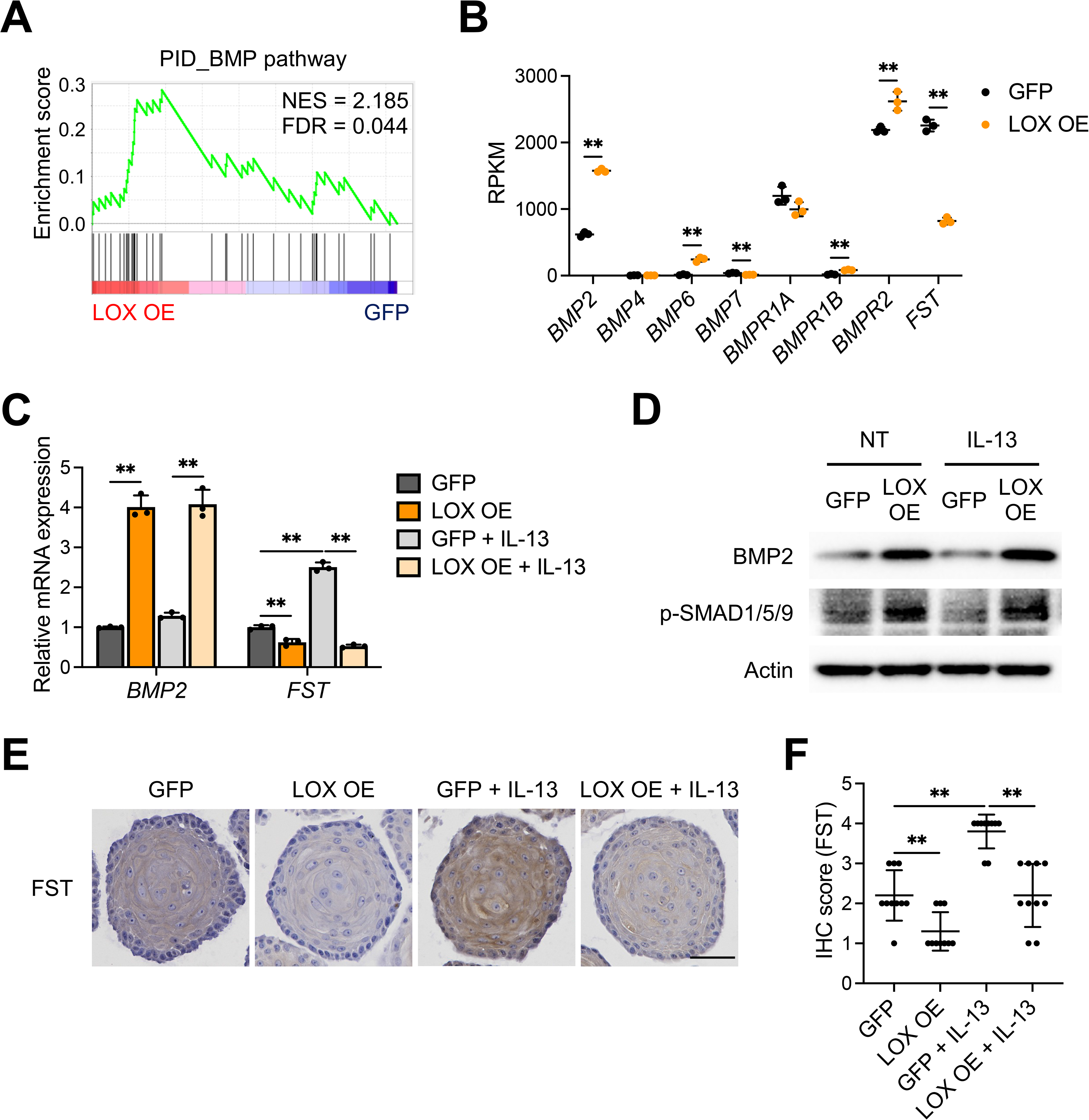
BMP signaling pathway is activated in LOX overexpressing organoids. (**A**) Gene Set Enrichment Analysis for BMP pathway in the Pathway Interaction Database (PID) based on the LOX overexpressing (LOX OE) organoids RNA sequencing data. (**B**) Expression of BMP and BMP receptor (BMPR) genes which relevant to the BMP pathway gene set, plotted as reads per kilobase per million (RPKM) (n = 3). (**C**-**F**) Validation of the BMP activation in EPC2-hTERT organoids. GFP and LOX OE organoids were cultured with or without IL-13 (10 μg/ml) from day 7 to 11 and then harvested at day 11. (**C**) Quantitative RT-PCR for *BMP2* and *FST* of the GFP and LOX OE organoids (n = 3). (**D**) Representative images of immunoblot for BMP2 and phospho-SMAD1/5/9 (p-SMAD1/5/9) and (**E**) immunohistochemistry staining for FST of the GFP and LOX OE organoids. Scale bar, 50 μm. (**F**) FST protein levels were quantified with the scoring described in the method (n = 10). Data are representative of three independent experiments and expressed as means ± SDs. Two-tailed Student’s t-test (**B**) and one-way analysis of variance (**C** and **F**) were performed for statistical analyses. \*\**P* < 0.01. FDR, false discovery rate; NES, normalized enrichment score; NT, nontreated.

We validated these findings by qRT-PCR, western blot, and immunohistochemistry in the setting of IL-13 stimulation (Figure 6C-F). Transcript levels of *BMP2* were increased by 4.0-fold and 3.2-fold in untreated and IL-13-stimulated LOX OE organoids compared to GFP organoids, respectively. IL-13 stimulus led to increased expression of *FST* in GFP organoids, while LOX overexpression reduced this effect by 0.62-fold and 0.21-fold in untreated and IL-13-stimulated LOX OE organoids, respectively (Figure 6C). Protein levels of BMP2 and downstream transcription factor phospho-SMAD1/5/9 (p-SMAD1/5/9) were also increased in LOX OE organoids (Figure 6D). FST, which was robustly expressed in IL-13 stimulated organoids, was decreased in the setting of LOX overexpression (Figure 6E and 6F). These data support a model in which overexpression of LOX inhibits FST leading to increased BMP signaling.

### BMP2 promotes cell differentiation in esophageal epithelium

Finally, we investigated whether BMP2 is involved in esophageal epithelial differentiation and barrier integrity. Treatment with recombinant BMP2 led to decreased *TP63* mRNA expression and increased *IVL*, *FLG*, *DSG1*, and *DSC1* mRNA expression in monolayer-cultured EPC2-hTERT cell (Figure 7A). Immunoblotting demonstrated similar results with increased p-SMAD1/5/9, IVL, and DSG1 in BMP2-treated cells (Figure 7B). Consistent with the results in monolayer culture, BMP2-treated organoids showed decreased expression of basal genes (SOX2 and TP63) and increased expression of differentiation and junctional protein genes (IVL, FLG, LOR, DSG1, and DSC1) (Figure 7C and 7D). BMP2 treatment also reduced organoid formation capacity, further supporting the notion that BMP2 enhances differentiation and reduces the basal population (Figure 7E). In summary, our findings suggest that LOX has a protective role in the esophageal epithelium in which it acts to restore homeostasis in EoE via activation of BMP signaling.

**Figure 7.**
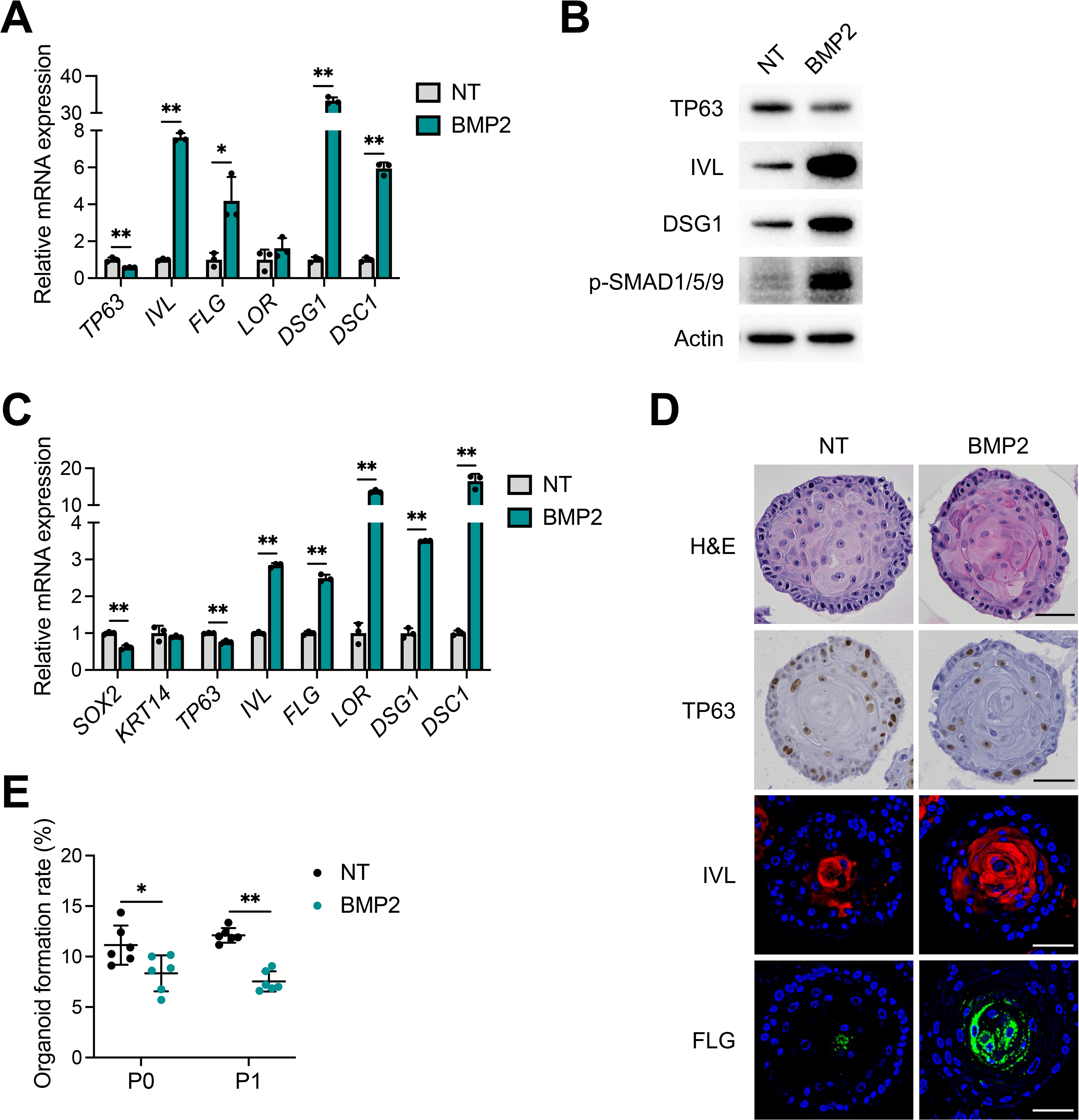
BMP2 treatment induces cell differentiation in esophageal epithelium. (**A** and **B**) BMP2 treatment in monolayer culture of EPC2-hTERT cells. EPC2-hTERT cells were treated with recombinant BMP2 protein (10 μg/ml) for 72 h in high-calcium (1.8 mM Ca^2+^) media. (**A**) Quantitative RT-PCR for *TP63*, *IVL*, *FLG*, *LOR*, *DSG1*, and *DSC1* in the EPC2-hTERT cells (n = 3). (**B**) Representative images of immunoblot for TP63, IVL, DSG1, and phospho-SMAD1/5/9 (p-SMAD1/5/9) of the EPC2-hTERT cells. (**C**-**E**) EPC2-hTERT organoids were treated with recombinant BMP2 protein (10 μg/ml) from day 7 to 11 and then harvested at day 11. (**C**) Quantitative RT-PCR for *SOX2*, *KRT14*, *TP63*, *IVL*, *FLG*, *LOR*, *DSG1*, and *DSC1* in the EPC2-hTERT organoids (n = 3). (**D**) Representative images of hematoxylin and eosin (H&E) staining, immunohistochemistry for TP63, and immunofluorescence staining for IVL (red) and FLG (green) of the EPC2-hTERT organoids. DAPI (blue). Scale bar, 50 μm. (**E**) Organoid formation rate (OFR) was assessed at day 11 (P0) and they were then passaged. OFR was assessed again at day 11 (P1). OFR was defined as the number of organoids (≥ 50 μm) divided by the total seeded cells (n = 6). Data are representative of three independent experiments and expressed as means ± SDs. Two-tailed Student’s t-test (**A**, **C**, and **E**) was performed for statistical analyses. \**P* < 0.05, \*\**P* < 0.01. NT, nontreated.

## Discussion

Lysyl oxidase is an extracellular matrix remodeling enzyme, which acts to cross link collagen thereby enhancing tissue stiffness. LOX is expressed in the epithelium of the prostate, retina, and skin as well as other locations (36–38), however its role in the esophageal epithelium in homeostasis and disease is unknown. Herein, we describe a non-canonical role for LOX in the esophageal epithelium and a potential protective role in epithelial differentiation and barrier integrity. Building upon our previous work demonstrating that LOX was upregulated in the EoE epithelium, we now show that LOX is expressed in the differentiated esophageal epithelium where it has non-collagen-crosslinking roles. LOX overexpression models demonstrate enhanced differentiation and barrier integrity even in the setting of IL-13 stimulation in the esophageal epithelium. Utilizing unbiased transcriptomic approaches, we found that the BMP pathway was enriched in LOX OE cells and that LOX OE cells have increased expression of BMP2, while BMP antagonist follistatin is decreased. This effect is maintained despite IL-13 stimulation. These results help to elucidate the contribution of LOX-supported differentiation and barrier integrity both in homeostasis and in the context of allergic insult.

Epithelial changes in EoE disrupt the mucosal barrier which normally provides protection from acid and food particles during normal swallows. Current treatment strategies are aimed at decreasing the invasion of inflammatory cells into the esophagus. However, we have found that epithelial changes persist even in patients in remission with low eosinophil counts (2). Thus, determining mechanisms to restore homeostasis to the esophageal epithelium represents an unexplored avenue of research with therapeutic potential. We now highlight a novel role for LOX in restoration of epithelial homeostasis. LOX silencing has been shown to impair keratinocyte differentiation of the skin (36). In fact, *in vitro* skin models show that LOX expression is increased in early differentiation and knockdown inhibits terminal differentiation. Taken together, these results suggest that increased LOX in the setting of inflammation may serve to re-establish differentiation and barrier during injury in the squamous mucosa.

Fibrosis is a major complication in EoE, and since LOX is a collagen crosslinker, it would be tempting to use LOX inhibitors to prevent fibrosis. However, our data demonstrating the protective roles of LOX in the esophageal epithelium in the context of Th2 inflammation would suggest that global LOX inhibition would have detrimental effects on the epithelial barrier. It would therefore be advantageous to target the crosslinker activity of LOX without effecting its non-crosslinking functions or vice versa. In that regard, our identification of the beneficial role of the LOX/BMP axis in the esophageal epithelium may represent a new approach to mitigating EoE, which could be employed via a topical approach. Topical steroid preparations are often employed in EoE to spare the negative consequences of systemic treatment with steroids (39,40). Furthermore, recombinant human BMP2 is currently approved by the Food and Drug Administration to promote bone healing in orthopedics (41–43). Building off of this and our *in vitro* findings of the protective role of BMP2 in the setting of IL-13 stimulation (Figure 7), it may be possible to employ preparations of BMP2 agonists to enhance esophageal epithelial barrier integrity.

BMP2, a member of the TGFβ superfamily, is essential for embryogenesis and development of the gut, and the BMP pathway affects morphogenesis of esophageal epithelial progenitor cells (44–47). We revealed that overexpression of LOX results in increased expression of BMP2 and increased squamous differentiation in esophageal organoids. Interestingly, downregulated BMP2 expression by fibroblast growth factor 9, which is upregulated in patients with EoE, has been proposed as the mechanism of hyperplasia in EoE (48). Furthermore, basal cell hyperplasia in the EoE epithelium, specifically in the setting of IL-13 stimulation, is associated with inhibition of the BMP pathway in human disease and murine models (34). Herein, we demonstrate the role of LOX specifically in differentiated cells to maintain barrier integrity and restore homeostasis. Future work will interrogate the mechanism by which LOX activates the BMP pathway to exert these protective roles. One possibility is downregulation of the BMP antagonist follistatin (49) by LOX. We demonstrated decreased follistatin levels in LOX OE organoids, even in the setting of IL-13 stimulation (Figure 6). Interestingly, IL-13/STAT6 pathway directly upregulates follistatin, and its increased levels have been reported in EoE patients and murine models (34). Correspondingly, knockdown of follistatin accelerates epithelial differentiation in esophageal cells. To date, although reactive oxidative stress has been shown to mediate the differentiation by the activated BMP in EoE (34), the specific mechanism remains to be elucidated.

One weakness of this study is that we rely on cell culture techniques as there is no *in vivo* model of LOX overexpression. LOX knock out mouse models would complement this work, however, these models are embryonic lethal and tissue specific knock out mouse strains do not exist. To contend with these limitations, we utilize 3D cultures (organoid and ALI culture) to replicate the proliferation and renewal patterns *in vitro*. Another potential weakness is that while LOX is upregulated in EoE, our model may be inducing higher overexpression levels than observed *in vivo*. However, the benefit of this model is that it allows for evaluation of the mechanistic effects of LOX without the secondary effects of inflammatory cytokines.

The esophageal epithelial barrier is maintained by an exquisitely regulated proliferation and differentiation gradient. The perturbations lead to symptoms as well as inflammation. Herein, we describe a novel role for LOX involving maintenance of differentiation and barrier integrity independent of its effects in subepithelial matrix remodeling. Epithelial LOX may serve to re-establish homeostasis in the setting of inflammation via BMP activation, underscoring the diverse functions of LOX in the esophagus. Investigation of the LOX-BMP pathway may provide novel insights into the pathogenesis and be a promising therapeutic approach for EoE. Further, elucidating these mechanisms may have implications for epithelial disruption in disorders beyond EoE, including caustic ingestions and gastroesophageal reflux disease.

## Methods

### Cell line and monolayer culture

EPC2-hTERT cells were cultured in keratinocyte-serum free medium with 0.09 mM Ca^2+^ (KSFM; Thermo Fisher Scientific Inc., Waltham, MA, USA) supplemented with bovine pituitary extract (50 μg/ml), human recombinant epidermal growth factor (1 ng/ml), and 1% penicillin-streptomycin. The cells were incubated at 37°C in a 5% humidified CO_2_ atmosphere (50,51).

### 3-dimensional esophageal organoid culture

EPC2-hTERT organoids were cultured as described previously (16–18). Briefly, EPC2-hTERT cells were dissociated into a single cell suspension and placed into Matrigel basement membrane matrix (Corning Inc., Corning, NY, USA) under KSFM modified with 0.6 mM Ca^2+^. Organoids were grown for 7 days, followed by treatment with 10 ng/ml IL-13, 10 ng/ml recombinant BMP2 protein (R&D Systems, Inc., Minneapolis, MN, USA), or vehicle (phosphate-buffered saline for IL-13 and hydrochloride for BMP2) for 4 days. We then used the day 11 organoids for further analyses. OFR was defined as the number of organoids ≥ 50 μm divided by the total seeded cells in each well (17,18).

### Air-liquid interface culture

EPC2-hTERT cells were seeded on transwell permeable supports with 0.4 μm pore (3470; Corning) and grown with KSMF (0.09 mM Ca^2+^) for initial 3 days to confluency. Cultures were then switched to high-calcium KSFM (1.8 mM Ca^2+^) for 5 days. The media was removed from the apical compartment on day 8 to induce epithelial differentiation and stratification. 10 μg/ml IL-13 (or vehicle) was applied in the basolateral compartment from day 9 to 14. In order to assess epithelial barrier integrity, TEER was measured with Epithelial Volt/Ohm (TEER) Meter (World Precision Instruments, Sarasota, FL, USA). The day 14 ALI-cultured epithelium was used for TEER measurement and histology.

### Bioinformatic analysis of single-cell RNA sequencing data

PDOs were stimulated with vehicle or IL-13 (10 μg/ml) from day 7 to day 11 and the day 11 PDOs were then dissociated into a single cell suspension for single-cell RNA sequencing (18). Dead Cell Removal Kit (Miltenyi Biotec, Bergisch Gladbach, Germany) was used to guarantee cell viability. The raw count matrix with barcode and feature information for each sample were imported and transformed to Seurat (version 4.2.0) objects for further processing. Genes expressed in 3 or fewer cells were excluded from analysis. To eliminate dead cells or doublets, cells with the expression of less than 700 or over 6000 genes, respectively, were excluded. Additionally, cells with over 10% of their transcripts consisting of mitochondrial genes were excluded. Seurat integration workflow was used to integrate the top 3000 variable genes, as anchors, across cells for control samples (52). After integration, dimensionality reduction used the genes and values that were pre-processed using the integration workflow. Principal component analysis was used for initial dimensionality reduction and later for clustering, resulting in 20 principal components. The components were then used as input to the UMAP dimensionality reduction procedure using 20 neighbors for local neighborhood approximation and embedding into 2 components for visualization. Clustering was initialized with a Shared Nearest Neighbor (SNN) graph by first determining 20 nearest neighbors for each cell, and then determined by a modularity optimization algorithm by Waltman and van Eck (53). Cell type annotations for each cluster are based on the expression of marker genes and DEGs. To better compare the IL-13 treated samples to the control data, IL-13 data were projected to control data after filtering and integration using Seurat projection/query workflow. For DEG analysis on the IL-13 treated PDOs, cells with a LOX expression greater than 0 were defined as LOX^+^ cells and cells with a LOX expression of 0 were defined as LOX^-^ cells in the superficial cluster.

### Trajectory analysis

Monocle 3 (version 1.0.0) was used to infer the trajectory analysis based on the single-cell RNA sequencing data. Seurat objects from upstream analysis were converted to Monocle objects, and then reversed graph embedding was applied to yield principal graph that is allowed to branch from the reduced dimension space (54). Pseudotime trajectory are defined and derived by selecting the specific cells as roots based on the prior knowledge of the cell type.

### Bulk mRNA sequencing and gene expression analysis

GFP and LOX OE organoids were grown for 11 days and then harvested for RNA sequencing. Sequencing libraries were constructed from total RNA (1 μg) using a TruSeq Stranded mRNA Library Prep (Illumina Inc., San Diego, CA, USA). RNA sequencing was performed on Illumina HiSeq2000 platform. We used kallisto (55) and human reference genome hg38 for alignment. Mapped reads were analyzed with DESeq2 (32). DEGs were determined as *P* < 0.05 and log_2_(fold change) ≥ 0.585 or ≤ -0.585. Using DEGs, Gene Ontology enrichment analysis was provided by topGO package (56). Top 5 enriched or depleted terms with the lowest FDR values in LOX OE organoids were selected. The ‘stat’ output field from DESeq2 was then used as input for GSEA (57,58) preranked analysis to identify enriched pathways from the PID.

### Lentivirus-mediated gene transfer

Lentiviral vectors pLX304-eGFP and pLX304-LOX were constructed by Gateway LR reaction of entry clones pENTR223-LOX (HsCD00378945, PlasmID Harvard Medical School) and pDONR221-eGFP (Addgene vector 25899) with destination vector pLX304 (Addgene vector 25890). VSV-g coated lentiviral particles were prepared by transfecting 293T cells with 9 μg of psPAX, 0.9 μg of pMD2.G, and 9 μg of overexpression vector using Lipofectamine 2000 Transfection Reagent (Thermo Fisher Scientific Inc.). EPC2-hTERT cells were transduced by spinfection and selected with 10 μg/ml blasticidin (59).

### Quantitative reverse transcription-polymerase chain reaction

RNA extraction and reverse transcription were carried out as described previously (21,51). Real-time qRT-PCR was performed with TaqMan Gene Expression Assays (Thermo Fisher Scientific Inc.) for *LOX* (Hs00942480_m1), *SOX2* (Hs01053049_s1), *KRT14* (Hs03044364_m1), *TP63* (Hs00978340_m1), *IVL* (Hs00846307_s1), *FLG* (Hs00856927_g1), *LOR* (Hs01894962_s1), *BMP2* (Hs00154192_m1), *FST* (Hs01121165_g1), and glyceraldehyde-3-phosphate dehydrogenase (*GAPDH*; Hs02786624_g1) using the StepOnePlus Real-Time PCR System (Thermo Fisher Scientific Inc.). Relative mRNA levels of each gene were normalized to *GAPDH* levels as a housekeeping control.

### Western Blot

Whole-cell lysates from cells in monolayer culture and 3D organoids were prepared as described previously (17). Equivalent amounts (20-40 μg) of protein were loaded into a NuPAGE 4 to 12% Bis-Tris gel. Following electrophoresis, transfer to a polyvinylidene difluoride membrane, and blocking with 5% bovine serum albumin or non-fat milk, membranes were incubated with primary antibodies at 4°C overnight. The primary antibodies used were follows: anti-LOX (1:500; NB100-2527; Novus Biologicals, Centennial, CO, USA), anti-TP63 (1:1000; ab124762; Abcam, Cambridge, UK), anti-IVL (1:1000; I9018; Millipore Sigma, Burlington, MA, USA), anti-DSG1 (1:1000; sc-137164; Santa Cruz Biotechnology, Dallas, TX, USA), anti-BMP2 (1:1000; ab214821; Abcam), anti-phospho-SMAD1/5/9 (1:1000;13820S; Cell Signaling Technology, Danvers, MA, USA), and anti-β-actin (1:5000; A5316; Millipore Sigma). Immunoblots were detected with an appropriate horseradish peroxidase (HRP)-conjugated secondary antibody (1:2000; NA934 or NA 931; Amersham BioSciences, Buckinghamshire, UK) by ECL detection (Bio-Rad Laboratories, Hercules, CA, USA). β-Actin served as a loading control.

### Immunohistochemistry and immunofluorescence

Organoids were fixed and embedded as described previously (16,17), and subjected to hematoxylin and eosin staining, immunohistochemistry, and immunofluorescence.

For immunohistochemistry, after deparaffinization and rehydration, antigens were retrieval by high-pressure cooking, Following, peroxidase quenching and blocking with an appropriate serum (Jackson ImmunoResearch), sections were incubated with primary anti-TP63 monoclonal antibody (1:1000; ab124762; Abcam) and anti-follistatin monoclonal antibody (1:50; MAB669; R&D Systems, Inc.) at 4°C overnight and then with an appropriate biotinylated secondary antibody (1:200; Vector Laboratories, Burlingame, CA, USA). The signal was detected with VECTASTAIN Elite ABC-HRP Kit (PK-6100; Vector Laboratories). DAB Substrate Kit (SK-4100; Vector Laboratories) was used for color reaction. For follistatin staining, slides were quantified with following score: 1 is negative to weak, 2 is weak to moderate, 3 is moderate to strong, and 4 is strong staining. Ten organoids per condition were used for the evaluation.

For immunofluorescence, sections were stained with primary anti-IVL monoclonal antibody (1:100, I9018; Millipore Sigma), anti-FLG monoclonal antibody (1:100; MA5-13440; Thermo Fisher Scientific Inc.), and anti-DSG1 monoclonal antibody (1:100; sc-137164; Santa Cruz Biotechnology) at 4°C overnight and then with Cy2 or Cy5-AffiniPure Donkey Anti-Mouse IgG (H+L) secondary antibody (1:500; 715-225-150 or 715-175-150; Jackson ImmunoResearch, West Grove, PA, USA) at room temperature for 1 h. Nuclei were stained with 4′,6-diamidino-2-phenylindole (DAPI; 17985-50; Electron Microscopy Sciences, Hatfield, PA, USA). Images were taken with an All-in-One Fluorescence Microscope BZ-X710 (KEYENCE Corp., Osaka, Japan). The images were evaluated in 5 different locations in high-power field per condition, and representatives are shown.

### Statistical analysis

Data are presented as means ± standard deviations (SDs). Continuous variables were analyzed by two-tailed Student’s t-test for two independent groups or analysis of variance for multigroup. All statistical analyses were conducted with GraphPad Prism (GraphPad Software, San Diego, CA, USA). A *P* value of less than 0.05 was considered statistically significant.

All authors had access to the study data and had reviewed and approved the final manuscript.

## Study approval

Not applicable.

## Acknowledgments

We thank to the Molecular Pathology and Imaging Core (Kate Bennett, Rebecca Ly, and Jonathan P Katz) for technical support. Schematics were created with BioRender.com.

---

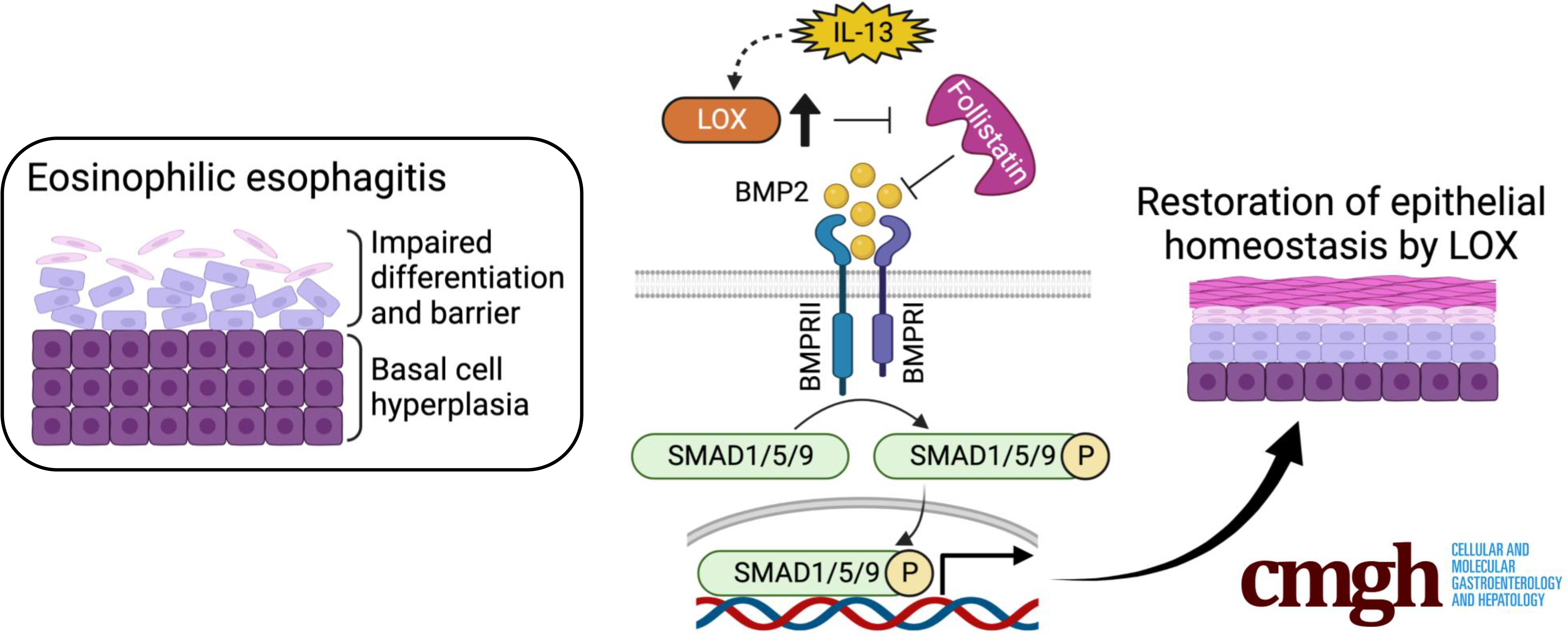

